# The history of body size evolution in termites

**DOI:** 10.1101/2021.09.30.462579

**Authors:** Nobuaki Mizumoto, Thomas Bourguignon

## Abstract

Termites are social cockroaches. Because non-termite cockroaches are larger than basal termite lineages, which themselves include large termite species, it has been proposed that termites experienced a unidirectional body size reduction since they evolved eusociality. However, the validity of this hypothesis remains untested in a phylogenetic framework. Here, we reconstructed termite body size evolution using head width measurements of 1638 modern and fossil termite species. We found that the unidirectional body size reduction model was only supported by analyses excluding fossil species. Analyses including fossil species suggested that body size diversified along with speciation events and estimated that the size of the common ancestor of modern termites was comparable to that of modern species. Our analyses further revealed that body size variability among species, but not body size reduction, is associated with features attributed to advanced termite societies. Our results suggest that miniaturization took place at the origin of termites, while subsequent complexification of termite societies did not lead to further body size reduction.

## Introduction

Body size diversification is an indicator of ecological diversification [1–3]. Diversification occurs when new resources or niches become available [4,5], often leading to the evolution of new phenotypes (i.e., key innovations [6–8]). The evolution of eusociality is a major evolutionary transition [9], which potentially marked the rise of several adaptive radiations and led to phenotypic diversification, including body size diversification. Social insects, especially ants and termites, are among the most successful and abundant terrestrial animals [10,11]. Their colonies typically contain many individuals belonging to distinct specialized phenotypic castes, which vary in size in a species-specific manner. However, the factors responsible for body size variation among species, and the role of social evolution, remain unclear. This problem can be addressed by analyses of body size measurements in a comparative phylogenetic framework.

Termites are a group of social insects that share a common ancestor with the wood roach *Cryptocercus*, from which they diverged some 170 million years ago [12–14]. The genus *Cryptocercus* contains only 12 of the 4,600 described cockroach species [15], while termites include over 3,000 described modern species [16,17], considerably varying in body size (Fig. 1). The unidirectional reduction in body size is believed to be a general evolutionary trend in termites [18]. Small body size enables more individuals to inhabit small pieces of wood, perhaps allowing larger and more complex societies to evolve [19]. Small body size also allows for nutrient economy, especially nitrogen, which is limiting in wood-boring insects [18]. The unidirectional body size reduction is supported by the observations that all species of *Cryptocercus* are much larger than species of termites (Fig. 1), and modern representatives of early-diverging termite lineages are generally large [18,20]. However, this idea has never been adequately tested and relies on comparisons among termite families and cockroaches. In addition, fossil species have not been taken into consideration, despite the existence of key fossil species with small body size, such as *Melqartitermes*, one of the oldest know termite fossils [21], and *Nanotermes*, the smallest termite species ever known to have existed [22].

**Fig. 1.**
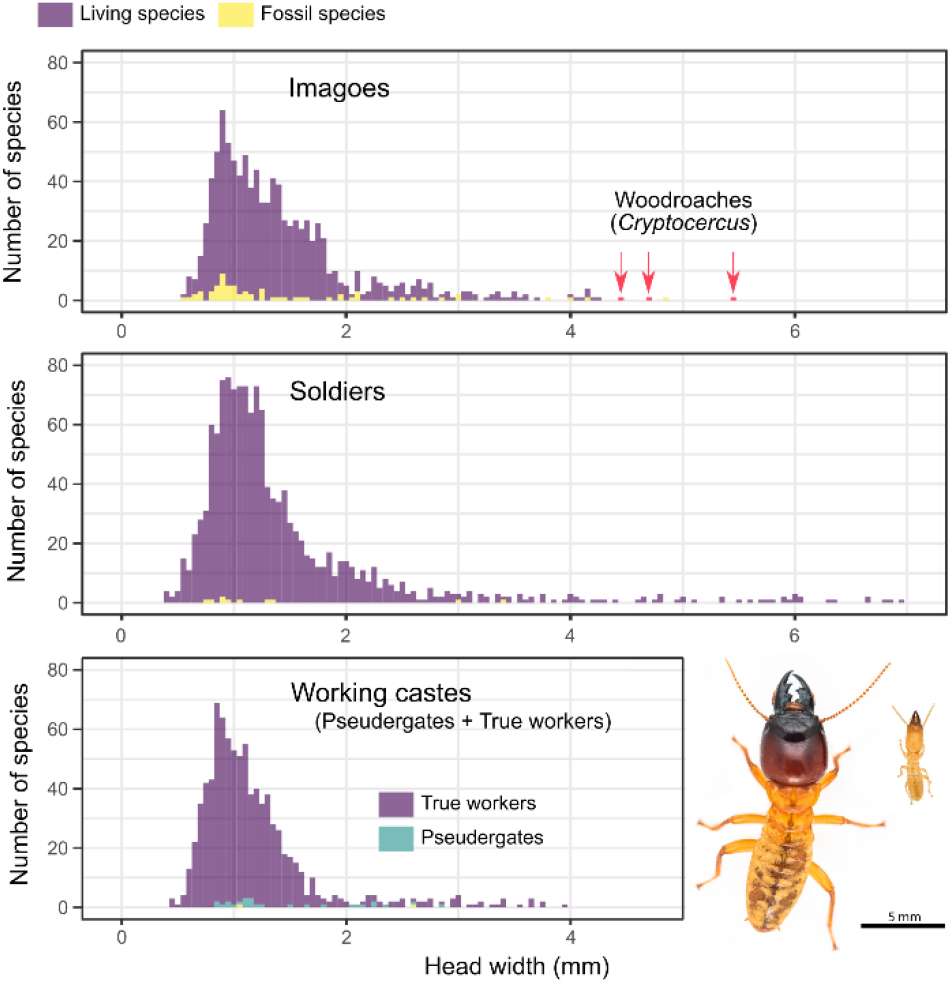
Distribution of head width across termite species. We used the data of the largest morph when within-caste polymorphism was present (i.e., major workers and major soldiers). The photos show the soldiers of a large species (*Hodotermopsis sjostedti*) and a small species (*Reticulitermes okinawanus*) (photos: Ales Bucek).

Another factor that has possibly affected body size evolution in termites is sociality. While all termites are eusocial, the level of social complexity varies among termite species and can be roughly approximated by nesting strategies and developmental pathways (Fig. 3A). Social complexity is presumably the lowest in one-piece nesting termites that nest in the piece of wood on which they feed, followed by multiple-piece nesting termites that feed on multiple wood items connected by networks of galleries, and is the highest in separate-piece nesting termites that build large nest structures separated from their food sources [23– 25] (Fig. 3A). The construction of large nests, separated from food sources, is undoubtedly a derived trait in termites [25,26], enabling colonies to settle over long periods. Developmental pathways are also variable among termite species, with two distinct types: the linear developmental pathway and the forked developmental pathway [27,28] (Fig. 3A). Species with a linear developmental pathway are often considered socially primitive as they lack a true worker caste. In these species, colony tasks are performed by immatures, called “pseudergates,” which retain the potential to develop into alate imagoes. In contrast, species with a forked developmental pathway possess a caste of “true workers” that irreversibly deviates from the imaginal developmental line at an early developmental stage and cannot molt into alate imagoes. Due to this additional caste, species with true workers have increased phenotypic and behavioral plasticity [29], potentially allowing for the evolution of more complex social systems. Whether a separate-piece nesting strategy and the presence of a true worker caste are linked to body size evolution remains unclear.

In this study, we reconstructed termite body size evolution using head width data collected from 153 papers (Data S1). We used head width as a proxy for body size because it has been consistently measured since the inception of termite taxonomy [18,30]. We fit seven evolutionary models on two phylogenetic trees, reconstructed with and without fossil species, to identify the most plausible scenario of body size evolution in termites. We also examined whether characteristics traditionally attributed to complex termite societies, including separate-piece nesting strategy, the presence of a true worker caste, and large colony size, are linked to body size evolution. Furthermore, we investigated how termite social evolution has shaped body size variation among castes, including body size variation among alate imagoes, soldiers, and working castes (pseudergates or true workers).

## Methods

### Data collection

We collected termite head width data from the literature. Most of the measurements were retrieved from taxonomic papers cited in the Termite Database [31]. Data were obtained for workers (or pseudergates), soldiers, and alate imagoes. We obtained one representative value for each caste and species. We used species mean head width values, mid-range value (calculated from the minimum and maximum values for the species for which only ranges were reported), or head width of the holotype, in this order of priority. For species displaying polymorphism within worker or soldier castes, we used the measurements from the largest subcaste (i.e., major workers and major soldiers). Workers and soldiers usually lack eyes, while eyes may be included or excluded from head width measurements of imagoes. In all cases, we used head width data of alate imagoes including eyes, and did not consider measurements that were explicitly taken without eyes. We assumed that eyes were included in head width measurements when the authors made no mention of eyes because “maximum width of head with eyes” has been the recommended measurement in termite taxonomy [30], and most studies include eyes in the measurements. The list of measurements and the source of these measurements are available in Data S1-2.

We classified termites based on their nesting habitats, the presence of true workers, and colony size. We recognized three categories of termite nesting habitats, as described by Abe (1987) [23]: one-piece, multiple-piece, and separate-piece nesters (Fig. 3A). One-piece nesters include species with nests consisting of a single piece of wood (including damp wood, dry wood, and dead branches on living trees), serving both as shelter and food source. One-piece nesters include *Zootermopsis*, all genera of Stolotermitidae, Stylotermitidae, Serritermitidae, almost all species of Kalotermitidae, *Prorhinotermes*, and *Termitogeton*. Multiple-piece nesters include species forming colonies encompassing multiple wood pieces among which individuals travel through belowground tunnels or aboveground shelter tubes. Multiple-piece nesters include *Mastotermes, Hodotermopsis, Paraneotermes*, and most species of Rhinotermitidae. Separate-piece nesters build nests physically separated from their food sources, which can be subterranean, built at the soil surface in the shape of large mounds, or arboreal. The separate-piece nesters include all Hodotermitidae and most species of Termitidae. Similarly, we classified termites into two categories based on the presence of true workers (Fig. 3A). *Mastotermes*, Hodotermitidae, Rhinotermitinae, *Reticulitermes, Heterotermes, Coptotermes*, and all Termitidae have true workers, while other termite lineages are devoid of a true worker caste and rely on pseudergates for colony tasks. We obtained maximum colony size estimates from one previous study [32]. Note that the methods used to infer colony size varied among studies and are prone to errors.

### Phylogeny

We used MRBAYES version 3.2.7 [33] to reconstruct a time-calibrated phylogenetic tree of extant and fossil termites. We used a relaxed clock model that combined molecular data of extant termite species and morphological characters of extant and fossil termite species. Our phylogenetic tree was composed of 183 taxa, including 138 modern termite species, each belonging to a distinct genus, 39 fossil termite species, and six outgroups, including *Cryptocercus*, four other cockroaches, and one mantis. The molecular data included 139 (133 termites + six outgroups) previously published mitochondrial genomes available on GenBank [13,34,43,35–42] (Data S3). All mitochondrial genomes were annotated using the MITOS webserver [44] with the invertebrate mitochondrial genetic code and default parameters. The two ribosomal RNA genes, 22 transfer RNA genes, and 13 protein-coding genes were aligned independently with MAFFT v7.300b using the options “--maxiterate 1000 –globalpair” for maximum accuracy [45]. Ribosomal RNA genes and transfer RNA genes were aligned as DNA sequences. Protein-coding genes were aligned as protein sequences and were back-translated to DNA sequences using pal2nal v14 [46]. We used the dataset published by [47] for the morphological data, which included 111 morphological characters scored across 82 taxa with five outgroups. We reduced the dataset to 79 taxa, including four outgroups. Modern taxa in this dataset were associated with one mitochondrial genome derived from the same species or another congeneric species when available. Because all modern species used in this study belonged to distinct genera, the morphological and molecular data always derived from specimens that were more closely related to each other than to any other taxa used for phylogenetic inferences. Fossil taxa were coded as “?” for molecular characters, and extant taxa without morphological data were coded as “?” for morphological characters. Our final dataset included 183 taxa, 145 of which were living taxa and 38 were fossil taxa. Both molecular and morphological data were available for 35 taxa, while 104 taxa and 44 taxa were exclusively represented by molecular data and morphological data, respectively. The molecular dataset was partitioned into four subsets: one combining the 12S and 16S rRNA genes, one combining the 22 tRNA genes, one for the first codon sites of protein-coding genes, and one for the second codon sites of protein-coding genes. The third codon position sites were excluded from the phylogenetic reconstruction because their substitution rate is too high to infer old divergences reliably. We used a GTR model with gamma-distributed rate variation across sites. The morphological data were assigned a MK+Γ model, with coding set to “variable” to account for acquisition bias [1,48]. To assist the analyses, we applied a series of topological constraints based on the topology reported in previous studies [13,14,47]. The nexus file, including the final alignment and the MrBayes block, and the reconstructed phylogenetic tree are available as supplementary material (Data S4, 5).

### Modeling of body size evolution

We combined our head width dataset with our phylogenetic tree to investigate various scenarios of termite body size evolution. We used imago head width as the representative body size measurements for each species because imago is the adult stage and the most common caste found in the fossil record. Taxa lacking head width data and taxa missing in our phylogenetic tree were not included in downstream analyses. The combination of the phylogenetic tree and head width data resulted in a phylogeny with 140 tips (113 modern genera + 27 fossil species, Fig. 2A) that we used for model fitting. We used genus-level information for modern termites because this taxonomic level is representative of the global evolutionary patterns found across termites. We used species-level data for fossils as fossils are scarce, and congeneric species sometime have different geological time. To account for the influence of within-genus variation on model fitting, we generated 100 sub-datasets by sampling one species for each genus at random. Every evolutionary model was fitted to every 100 sub-dataset.

**Fig. 2.**
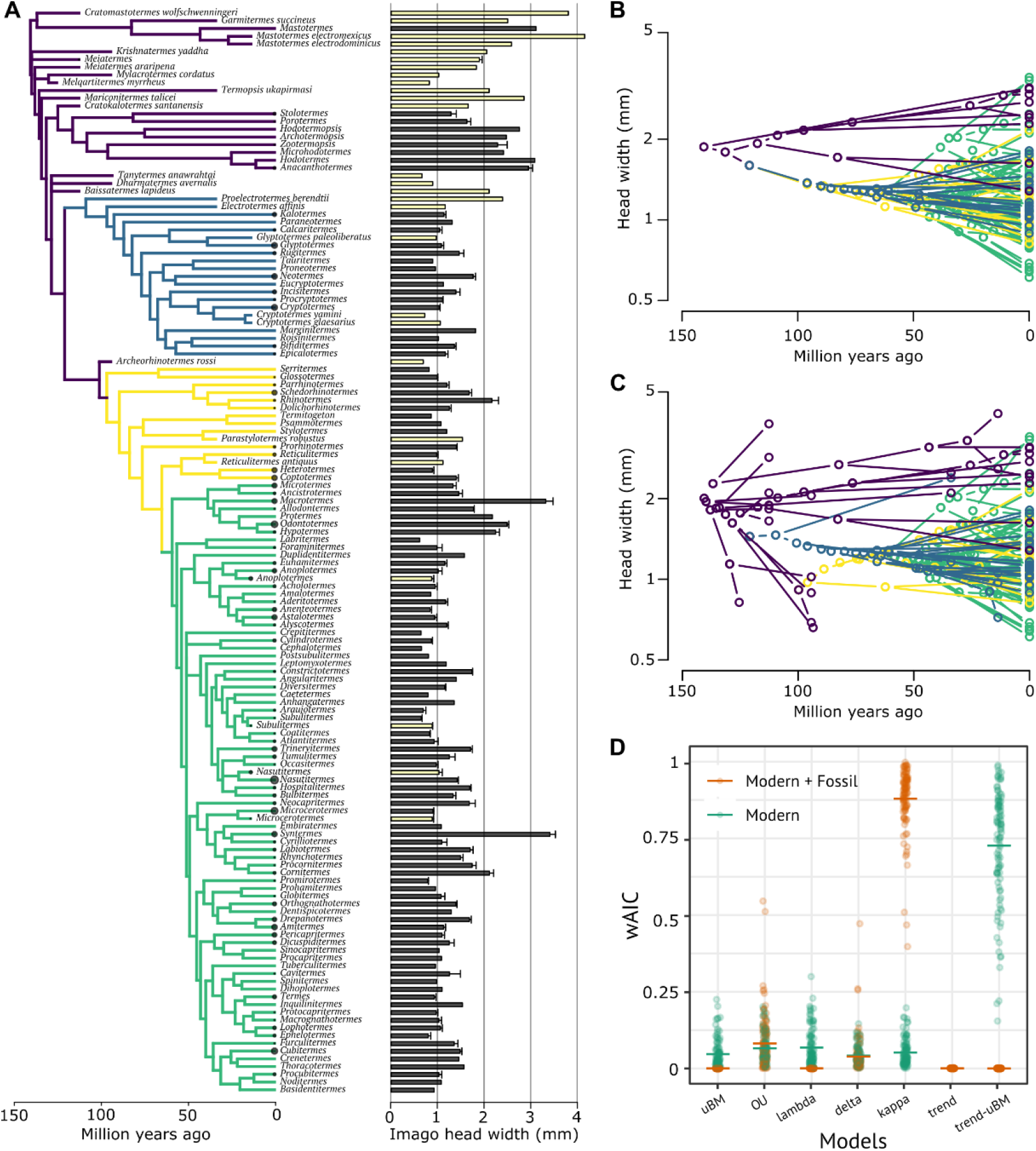
Evolution of termite imago head width. (*A*) Phylogenetic tree and head width data used for ancestral state reconstruction and model fitting. For modern genera, the barplot indicates the mean head width (estimated from all species composing each genus) and its associated standard error. For fossil species, we used species-level data when available. Filled bars indicate modern taxa, while open bars indicate fossil taxa. Circles at the tips of the phylogenetic tree represent sample size. (*B-C*) Traitgrams projecting the phylogeny and the evolution of head width in (*B*) modern termite genera and (*C*) modern and fossil termites combined. The traitgrams were generated using the function phenogram() of the R package phytools [63]. (*D*) Akaike weights for seven models fitted on the trees with modern genera only (green) and with both modern genera and fossil species (orange). For model fitting, we generated 100 datasets by subsampling the measurement of one species for each genus at random. The horizontal lines show the mean values.

We used the maximum likelihood method to fit process-based models of trait evolution on the head width data. We used the models supported by the fitContinuous() function of the R package geiger [49], including unbiased Brownian motion (BM), Brownian motion with a directional trend (trend), single optimum Ornstein-Urlenbeck process (OU), lambda, kappa, and delta. Note that we did not use the early burst model, which is also available in the geiger package, because it assumes a similar evolutionary process to delta model and cannot be applied to non-ultrametric trees with upward of 100 terminal branches [50]. We also implemented a model that explicitly describes an evolutionary scenario with a unidirectional decrease in body size in lower termites and no directional trend in higher termites (mixed model of trend and Brownian motion: trend-BM). We added this latter model for two reasons. First, higher termites are a derived group and have the largest body size diversity among termite families. Higher termites include many large species nested within lineages composed of species with small body sizes, which is in direct contradiction with a unidirectional body size reduction model. Therefore, the unidirectional body size reduction model appears to be invalid for higher termites but could be valid for lower termites. Second, a simple unidirectional trend model cannot fit datasets composed exclusively of modern species (see e.g., [51]). We compared the fit of these seven candidate models to (i) the dataset including both modern and fossil species and to (ii) the dataset including only modern species. The support of each model was compared using Akaike weights computed from AICc. We took the natural log values of head width data for model fitting. All analyses were performed using R v.4.0.1 [52]. The R scripts used in this study are available as supplementary material (Data S6).

### Phylogenetic comparison

The relationship between head width and social complexity was investigated with Bartlett’s test of homogeneity of variance and with phylogenetic generalized least squares (PGLS). Three biological traits were used as a proxy for social complexity: the presence of true workers, the nesting types, and the colony size. The Bartlett’s test was performed on the full head width measurement dataset containing species-level information because it did not account for the phylogenetic relationships among taxa. PGLS was performed on the head width measurement dataset summarized at the genus level and on the genus-level phylogenetic tree described above. We used mean head width values for each genus. We used the function pgls() of the R package caper_1.0.1 [53]. We carried out the analyses with Brownian, lambda, kappa, and delta models and used the best fit model identified by AIC. The presence of true workers, nesting types, or colony size was treated as fixed effects. As the presence of true workers, nesting types, and colony size are unknown for fossil species, we only used data obtained from modern species.

We also examined whether size variability among castes was affected by the presence of true workers. We calculated the proportional head width disparity [54] for the pairs imago-worker and imago-soldier using the following equations: (*Imago head width − Worker head width*) /*Worker head width* and (*Imago head width − Soldier head width*) /*Soldier head width*, respectively. To compare these disparity metrics, we used Bartlett’s test and PGLS as described above. We ran PGLS analyses twice, with true workers or pseudergates being the base level, to examine whether level means are significantly different from 0. We also carried out pairwise correlations of head width among castes using PGLS.

## Results and Discussion

### Evolution of head width in termites

Using literature data, we compiled a dataset including head width measurements of 1638 termite species. The dataset included 1562 modern termite species (911 species for imagoes, 1303 species for soldiers, 840 species for true workers, and 26 species for pseudergates) and 76 fossil species (69 species for imagoes, ten species for soldiers, and two species for workers). This dataset comprised nearly half of the described termite species, belonging to 287 genera from all families and subfamilies. The size of this dataset exceeded previous datasets of termite body size by more than one order of magnitude [18,55,56]. Head width ranged from 0.550 mm to 4.840 mm in imagoes, 0.385 mm to 6.960 mm in soldiers, and 0.459 mm to 3.975 mm in workers (Fig. 1). The species head width distribution was right-skewed (Fig. 1), as is the case in many other groups of insects [57,58].

By fitting various evolutionary models for termite body size evolution, we found that the inclusion of fossil species had large effects on the results. For the dataset composed of modern genera only, the best-fit model was a mixed model of trend and Brownian motion (trend-BM), which posits a unidirectional body size reduction in lower termites and non-directional size diversification in higher termites (Akaike weight; mean ± SD = 0.73 ± 0.19; estimated parameters in Table 1; Fig. 2B,D). These results are in line with the traditional view of a unidirectional decrease in body size, at least in lower termites. However, our analyses on the dataset, including both modern and fossil taxa, supported a different scenario for termite body size evolution (Fig. 2A,C). Model fitting showed that the kappa model best explained the imago head width evolution (Akaike weight; mean ± SD = 0.88 ± 0.10; estimated parameters in Table 1; Fig. 2D). The parameter κ < 1 indicates that the degree of body size divergence is associated with the number of cladogenetic events [59]. Notably, models representing a unidirectional decrease in body size (trend or trend-BM model) poorly fitted the dataset with fossils included (Akaike weight < 0.001; Table 1, Fig. 2D). These results reflect the existence of several fossil species with small head widths (e.g., *Melqartitermes, Mylacrotermes*, and *Tanytermes*) allied to basal termite lineages, contrasting with the modern early-diverging lineage representatives that are large species (Fig 2 A,C). These results highlight the importance of fossil inclusion for an accurate estimation of trait evolution [51,60].

**Table 1.** Results of the maximum likelihood model fitting. The representative parameter values are given for the analyses with and without fossils. For the Ornstein-Uhlenbeck model, the parameter is α; for the lambda model, the parameter is λ; for the delta model, the parameter is δ; for the kappa model, the parameter is κ. For the trend-uBM model, the parameter is mu, which was estimated from lower termite data. The best-fit model for each dataset is indicated in bold.

| Data | Model | AICc | wAIC | Root | Parameter |
| --- | --- | --- | --- | --- | --- |
| Modern | Brownian Motion | -116.11 (10.13) | 0.047 (0.047) | 1.69 (0.03) | 4.3e-04 (3.8e-05) |
|  | Ornstein Uhlenbeck | -117.29 (9.00) | 0.066 (0.047) | 1.58 (0.04) | 0.0067 (0.0017) |
|  | lambda | -117.02 (8.78) | 0.068 (0.062) | 1.68 (0.03) | 0.89 (0.054) |
|  | delta | -116.32 (9.24) | 0.041 (0.031) | 1.55 (0.04) | 1.9 (0.31) |
|  | kappa | -116.62 (9.92) | 0.052 (0.043) | 1.87 (0.06) | 0.64 (0.079) |
|  | **Trend-uBM** | **-122.58 (8.90)** | **0.73 (0.19)** | **1.84 (0.13)** | **-2.5e-04 (2.1e-04)** |
| Modern+Fossil | Brownian Motion | -111.43 (7.62) | < 0.001 | 1.76 (0.01) | 6.0e-04 (3.2e-05) |
|  | Ornstein Uhlenbeck | -122.46 (6.85) | 0.081 (0.09) | 1.73 (0.02) | 0.01 (7.6e-04) |
|  | lambda | -112.31 (7.14) | < 0.001 | 1.76 (0.01) | 0.97 (0.009) |
|  | delta | -120.99 (9.73) | 0.038 (0.061) | 1.97 (0.02) | 0.55 (0.036) |
|  | **kappa** | **-128.44 (8.92)** | **0.88 (0.1)** | **2.31 (0.04)** | **0.2 (0.051)** |
|  | Trend | -112.29 (7.73) | < 0.001 | 1.59 (0.02) | 0.0012 (1.1e-04) |
|  | Trend-uBM | -109.81 (8.34) | < 0.001 | 1.61 (0.01) | 0.001 (6.2e-05) |

All models invariably estimated smaller head width for the last common ancestor of modern termites than the head width of the wood roach *Cryptocercus*, the sister group of termites. The kappa model run on the dataset comprising modern and fossil taxa estimated the head width of the last common ancestor of termites at 2.31 ± 0.04 mm (Table 1). Although 91% of modern termite species are smaller (Fig. 1), this estimation is half the size of *Cryptocercus*, whose head width is nearly 5 mm (Fig. 1). These results indicate that body size reduction occurred conjointly with the evolution of eusociality in termites over the 20 million year period following the divergence of termites from *Cryptocercus* and preceding the origin of the last common ancestor of modern termites [14]. The alternative scenario is that the common ancestor of termites and *Cryptocercus* had a small body size, which implies the subsequent acquisition of larger body size by modern *Cryptocercus*. However, this scenario is unlikely given the comparatively large body size of other wood-feeding cockroach lineages, such as *Salganea* and *Panesthia* [61]. Consequently, the selection pressures acting on small body size were strong at the very origin of termites [18,62] and weakened after the birth of the most recent common ancestor of modern termites, with body size following a diversification process.

Our comparative analyses suggested no connection between average body size and traits considered to be linked to advanced sociality in termites (Fig. 3). After accounting for the phylogenetic relationship among genera, we found no significant correlation between imago head width and colony size (PGLS, λ, κ, δ transformation, *F*_1_ = 1.28, *P* = 0.27; Fig. 3D). Similarly, average imago head width was independent of the presence of a true worker caste (PGLS, λ, κ, δ transformation, *F*_1_ = 2.884, *P* = 0.09; Fig. 3B) and of the type of nesting strategy (PGLS, λ, κ, δ transformation, *F*_2_ = 2.18, *P* = 0.12; Fig. 3C). Thus, caste systems and nesting strategies have no influence on average body size evolution in termites. These results are consistent with the observations that body size is not predictive of social behavior [64,65]. However, we found that interspecific variation in body size was the largest in species with traits deemed socially advanced. The interspecific variance of imago head width was significantly higher in termites with a true worker caste (Bartlett test; *K*_1_ = 53.88, *P* < 0.001; Fig. 3B) (variance = 0.42) than in those without true workers (variance = 0.15). Similarly, the interspecific variance of imago head width was significantly different in termites with different nesting strategies (Bartlett test; *K*_2_ = 70.66, *P* < 0.001; Fig. 3C). The variance significantly increased (Bartlett test; *P* < 0.01 for all pairwise comparisons) along the following sequence: one-piece nesters (variance = 0.13), multiple-piece nesters (variance = 0.24), and separate-piece nesters (variance = 0.44). These results indicate that the characteristics of socially complex termite societies do not influence average termite body size but are linked to the emergence of more extreme body size.

**Fig. 3.**
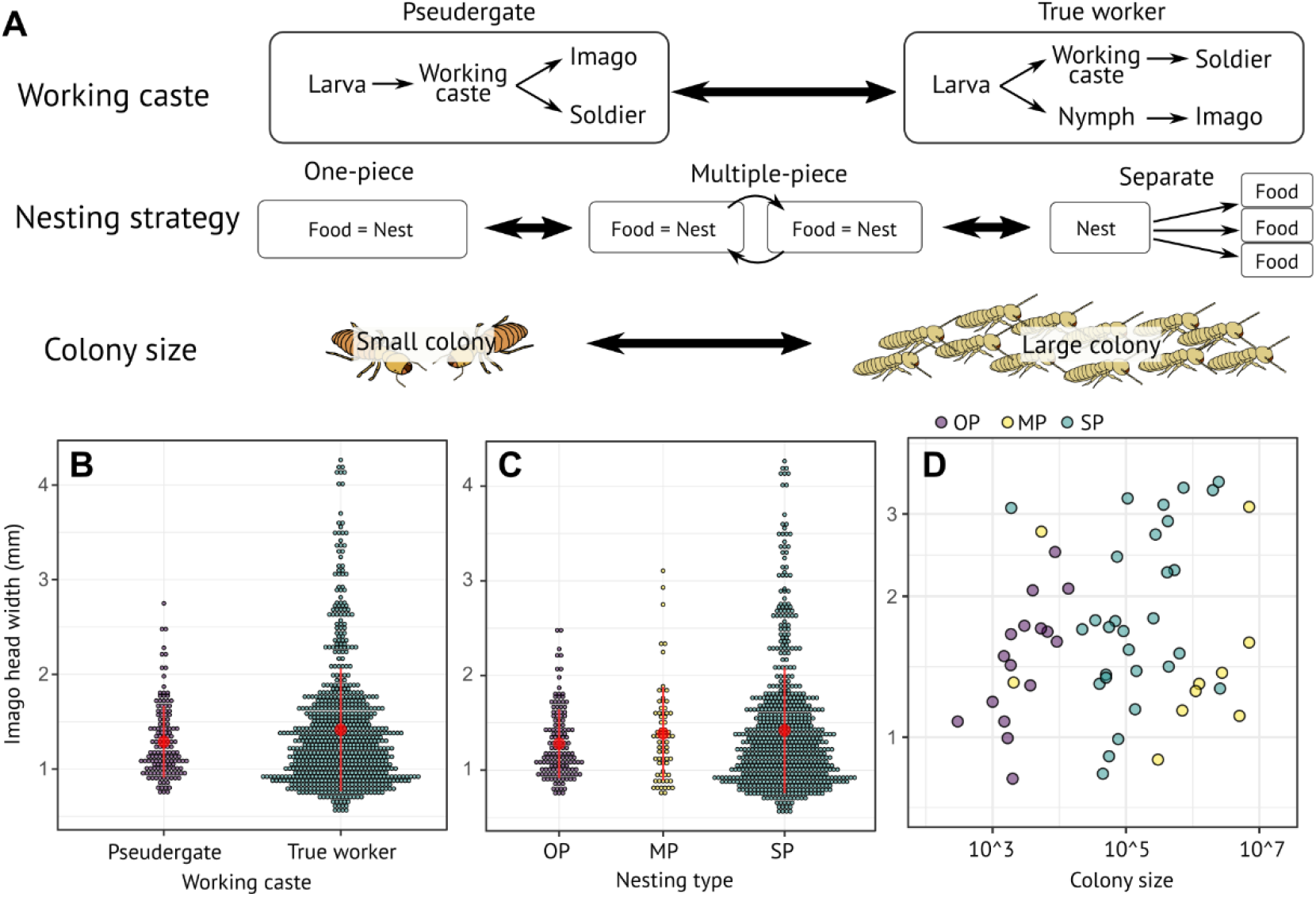
Relationship between social evolution and imago body size in termites. (A) Factors related to social evolution in termites: working caste, nesting strategy, and colony size. Some species lack true workers, and instead have pseudegates that retain the ability to differentiate in alate imagoes. In contrast, true workers are unable to molt into alate imagoes. Three nesting strategies are recognized: one-piece nesters, multiple-piece nesters, and separate-piece nesters. (B-C) Comparison of body size across groups with different levels of sociality. Red circles indicate the mean value, and red vertical lines are standard deviations. (C) OP, MP, and SP indicate one-piece nesters, multiple-piece nesters, and separate-piece nesters, respectively. (D) Relationship between colony size and imago head width.

### Differentiation of head width among castes

Division of labor among castes is the hallmark of social insects. To determine how social evolution affects termite body size evolution, we compared the head width of castes across termite genera. The head width of imagoes, soldiers, and the working castes strongly correlated to each other across genera (Fig.S1; PGLS, *P* < 0.001). However, the degree of correlation was dependent on the developmental pathway. Because the developmental lines of alate imagoes and workers diverged early on in the development of genera with true workers, and because pseudergates can differentiate into alate imagoes through a few molts [27] (Fig. 3A), we expected true workers to have a higher potential for phenotypic diversification. Our comparison between species with pseudergates and true workers found that imago head width was larger than pseudergate head width in 52.38% of examined species (11/21), while imago head width was larger than true worker head width in 85.65% of examined species (400/467). In some species with true workers, the imago head width was more than twice that of worker head width (e.g., *Microtermes*) (Fig. S1). The variance of head width disparity between imagoes and true workers was significantly greater than the variance of head width disparity between imagoes and pseudergates (Bartlett test, *K*_1_ = 4.677, *P* = 0.031; Fig. 4B). After accounting for the phylogenetic relationship among taxa, we found that the mean head width disparity between imagoes and working castes was not significantly different between genera with true workers and genera with pseudergates (PGLS, λ, κ, δ transformation, *F*_1_ = 3.22, *P* = 0.08; Fig. 4B), but the mean head width disparity was significantly greater than 0 in genera with true workers (PGLS, intercept, true workers: *t* = 2.858, *P* = 0.005, pseudergates: *t* = 0.797, *P* = 0.428; Fig. 4B). Therefore, the presence of a true worker caste allows alate imagoes to grow substantially larger than workers and is associated with a greater variation of head width disparity between alate imagoes and working castes. This link between developmental pathway and size diversity is paralleled in ants [66], suggesting that it represents a common characteristic of body size evolution in social insects.

**Fig. 4.**
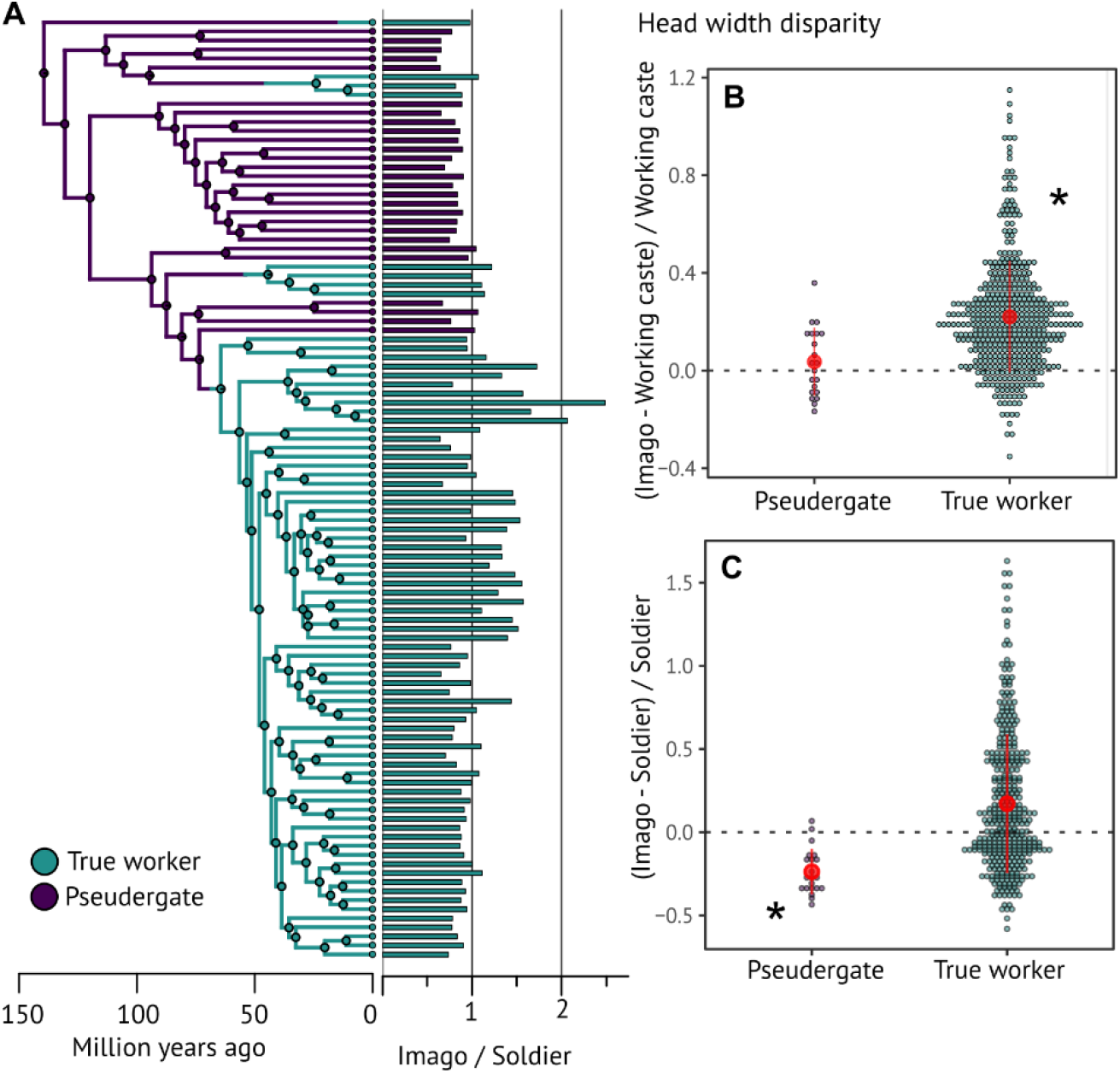
Disparity of head width among castes across the phylogenetic tree of termites. (A) Ancestral state reconstruction of true workers with head width disparity among castes. The reconstruction was carried out on a genus-level phylogeny, including taxa with data on both imago and soldier head width. Posterior probabilities of ancestral states were estimated using 100 stochastic mappings with the function make.simmap() of the R package phytools 0.7-47 [63]. An example of stochastic mappings is shown for the ancestral state reconstruction of each ratio. (B) Comparison of imago-working caste head width disparity for genera with pseudergates and true workers. (C) Comparison of imago-soldier head width disparity for genera with pseudergates and true workers. Red dots indicate the mean, and red vertical lines indicate the standard deviation. Asterisks denote that the mean of head width disparity is significantly different from 0, which indicates that imago head width is larger than true worker head width, or that soldier head width is larger than imago head width (PGLS; *P* < 0.05).

We carried out the same analyses on imago head width and soldier head width. Because all soldiers differentiate from the working castes, we expected soldiers to have a higher potential for phenotypic diversification in taxa with true workers, as we found was the case for the working castes. We found that most species with pseudergates had larger soldiers than imagoes (92.25%, 119/129), while soldiers were larger than imagoes in fewer species with true workers (46.13%, 280/607). The interspecific variation of head width disparity between imagoes and soldiers was significantly larger in termites with true workers than in termites with pseudergates (Bartlett test, *K*_2_ = 14.13, *P* < 0.001; Fig. 4C). After accounting for the phylogenetic relationship among taxa, we found that the mean head width disparity between imagoes and soldiers was larger in genera with true workers than in genera with pseudegates (PGLS, λ, κ, δ transformation, *F*_2_ = 11.32, *P* = 0.001; Fig. 4C), and that soldiers were in average larger than imagoes in genera with pseudergates, but not in genera with true workers (PGLS, intercept, with pseudergates: *t* = -4.790, *P* < 0.001, with true workers: *t* = -0.255, *P* = 0.799; Fig. 4C). The tendency of soldiers to be larger in species with pseudergates is linked to their one-piece nesting strategies. One-piece nesters are generally defended by phragmotic soldiers that plug galleries with their heads, which, to be efficient, need to have larger heads than other colony members. In contrast, the colonies of multiple-piece nesters and separate-piece nesters extend across larger areas and rely on soldiers employing diverse defensive strategies [67,68]. The greater variation of head width disparity between alate imagoes and soldiers reflects this diversity of defensive strategies, as to be optimal, each strategy required soldiers of different sizes.

## Conclusions

Termites have smaller body sizes than other wood-feeding cockroaches. Unidirectional body size reduction was believed to be the process behind termite body size evolution, at least in lower termites. However, we found that the unidirectional body size reduction hypothesis is only supported when fossil species are excluded from the analyses. Phylogenetic analyses including fossil species indicate that body size evolution was not a unidirectional process. Instead, body size reduction preceded the origin of the last common ancestor of modern termites, which already possessed a smaller body size than cockroaches. Thereafter, termite body size diversified along with cladogenetic events. Interestingly, a similar pattern was observed for the head width evolution of turtle ants [54]. Our results suggest that the body size range among early termite species was relatively similar to that found in modern termite species.

The acquisition of a diet based on wood, which is uncommon among animals, likely had a modest impact on termite body size diversification, as suggested by the apparent absence of body size reduction and diversification in the wood-feeding *Cryptocercus*, the sister group of termites. In contrast, the evolution of eusociality may have been one factor that promoted body size diversification in termites. Notably, termite body size diversification is linked to nesting strategies and developmental pathways, two biological traits closely associated with social complexity. Variation in body size among termite species was greater in taxa possessing a true worker caste and in separate-piece nesters, indicating that social complexity increases body size variation (Fig. 3).

Body size scales with various traits, including metabolism, abundance, and movements [57,69,70], which can also mediate social interactions between individuals. However, although many models described the interspecific variation of collective building in social insects [71–75], body size has rarely been implemented as a parameter. Further studies are needed to explore how species with considerable body size differences can build nests of similar size (e.g., *Macrotermes* and *Nasutitermes* build large mounds with the former having head width twice as large as the latter). Also, because closely related species often have similar body sizes, body size is often a confounding variable of other physiological and behavioral traits. To account for the effects of body size, the analysis of large body size dataset alongside phylogeny is essential. Our study paves the way for future comparative studies that aim to shed light on the ecology and evolution of animal society.

## Acknowledgments

We thank Christine Nalepa for providing head width data; Jigyasa Arora, Ales Bucek, Simon Hellemans, Yukihiro Kinjo, and Menglin Wang for help during the data analysis; Ales Bucek for photographs; Tomonari Nozaki and Hiroyuki Shimoji for helpful comments on the data; the members of the Evolutionary Genomics Units at OIST for inspiring scientific discussions. N.M. thanks his wife and son for allowing him to manually create a termite head width database during COVID-19 pandemic home quarantine. This study was supported by a JSPS Research Fellowship for Young Scientists, SPD and CPD (20J00660) to N.M., and OIST core funding.

## Author contributions

Nobuaki Mizumoto: Conceptualization, Data Curation, Formal Analysis, Funding Acquisition, Investigation, Methodology, Resources, Validation, Visualization, Writing-Original Draft Preparation. Thomas Bourguignon: Conceptualization, Data Curation, Project Administration, Resources, Supervision, Writing-Review & Editing.

## References

1. Slater GJ. 2013 Phylogenetic evidence for a shift in the mode of mammalian body size evolution at the Cretaceous-Palaeogene boundary. Methods Ecol. Evol. 4, 734–744. (doi:10.1111/2041-210X.12084)

2. Harmon LJ et al. 2010 Early bursts of body size and shape evolution are rare in comparative data. Evolution (N. Y). 64, 2385–2396. (doi:10.1111/j.1558-5646.2010.01025.x)

3. Erwin DH. 2015 Novelty and innovation in the history of life. Curr. Biol. 25, R930–R940. (doi:10.1016/j.cub.2015.08.019)

4. Schluter D. 2000 The ecology of adaptive radiation. Oxford: Oxford University Press.

5. Stroud JT, Losos JB. 2016 Ecological Opportunity and Adaptive Radiation. Annu. Rev. Ecol. Evol. Syst. 47, 507–532. (doi:10.1146/annurev-ecolsys-121415-032254)

6. Simpson GG. 1953 The Major Features of Evolution. New York: Columbia Univ. Press.

7. Rabosky DL. 2017 Phylogenetic tests for evolutionary innovation: The problematic link between key innovations and exceptional diversification. Philos. Trans. R. Soc. B Biol. Sci. 372. (doi:10.1098/rstb.2016.0417)

8. Hunter JP. 1998 Key innovation and ecology of macroevolution. Trends Ecol. Evol. 13, 31–36.

9. Smith JM, Szathmary E. 1997 The Major Transitions in Evolution. Oxford University Press.

10. Tuma J, Eggleton P, Fayle TM. 2019 Ant-termite interactions: an important but under-explored ecological linkage. Biol. Rev., brv.12577. (doi:10.1111/brv.12577)

11. Wilson EO, Hölldobler B. 2005 Eusociality: Origin and consequences. Proc. Natl. Acad. Sci. U. S. A. 102, 13367–13371. (doi:10.1073/pnas.0505858102)

12. Lo N, Tokuda G, Watanabe H, Rose H, Slaytor M, Maekawa K, Bandi C, Noda H. 2000 Evidence from multiple gene seqeunces indicates that termites evolved from wood-feeding cockroaches. Curr. Biol. 10, 801–804. (doi:10.1016/S0960-9822(00)00561-3)

13. Bucek A, Šobotník J, He S, Shi M, McMahon DP, Holmes EC, Roisin Y, Lo N, Bourguignon T. 2019 Evolution of termite symbiosis informed by transcriptome-based phylogenies. Curr. Biol. 29, 3728-3734.e4. (doi:10.1016/j.cub.2019.08.076)

14. Bourguignon T et al. 2015 The evolutionary history of termites as inferred from 66 mitochondrial genomes. Mol. Biol. Evol. 32, 406–421. (doi:10.1093/molbev/msu308)

15. Beccaloni GW. 2014 Cockroach species file online. Version 5.0/5.0. World Wide Web Electron. Publ.

16. Arumugam N, Mohd Kori NS, Rahman H. 2018 Termites identification. In Termites and Sustainable Management, pp. 27–45. Springer International Publishing. (doi:10.1007/978-3-319-72110-1_2)

17. Krishna K, Grimaldi DA, Krishna V, Engel MS. 2013 Treatise on the Isoptera of the World. Bull. Am. Museum Nat. Hist. 377, 1989–2433. (doi:10.1206/377.6)

18. Nalepa CA. 2011 Body size and termite evolution. Evol. Biol. 38, 243–257. (doi:10.1007/s11692-011-9121-z)

19. Korb J. 2008 The Ecology of Social Evolution in Termites. In Ecology of Social Evolution, pp. 151–174. Springer Berlin Heidelberg. (doi:10.1007/978-3-540-75957-7_7)

20. Grimaldi DA, Engel MS, Krishna K. 2008 The species of Isoptera (insecta) from the Early Cretaceous Crato Formation: A revision. Am. Museum Novit. 3626, 1–30. (doi:10.1206/616.1)

21. Engel MS, Grimald DA, Krishna K. 2007 Primitive termites from the Early Cretaceous of Asia (Isoptera). Staatl. Museum für Naturkd. 371, 1–32.

22. Engel MS, Grimald DA, Nascimbene PC, Singh H. 2011 The termites of Early Eocene Cambay amber, with the earliest record of the Termitidae (Isoptera). Zookeys 148, 105–123. (doi:10.3897/zookeys.148.1797)

23. Abe T. 1987 Evolution of life types in termites. In Evolution and Coadaptation in Biotic Communities (eds S Kawano, J Connell, T Hidaka), pp. 125–148. Tokyo: University of Tokyo Press.

24. Emerson EA. 1938 Termite nests: a study of the phylogeny of behavior. Ecol. Monogr. 8, 247–284.

25. Mizumoto N, Bourguignon T. 2020 Modern termites inherited the potential of collective construction from their common ancestor. Ecol. Evol., ece3.6381. (doi:10.1002/ece3.6381)

26. Inward DJG, Vogler AP, Eggleton P. 2007 A comprehensive phylogenetic analysis of termites (Isoptera) illuminates key aspects of their evolutionary biology. Mol. Phylogenet. Evol. 44, 953–967. (doi:10.1016/j.ympev.2007.05.014)

27. Roisin Y. 2000 Diversity and evolution of caste patterns. In Termites: Evolution, Sociality, Symbioses, Ecology, pp. 95–119. Springer Netherlands. (doi:10.1007/978-94-017-3223-9_5)

28. Korb J, Hartfelder K. 2008 Life history and development - A framework for understanding developmental plasticity in lower termites. Biol. Rev. 83, 295–313. (doi:10.1111/j.1469-185X.2008.00044.x)

29. Nozaki T, Matsuura K. 2019 Evolutionary relationship of fat body endoreduplication and queen fecundity in termites. Ecol. Evol. 9, 11684–11694. (doi:10.1002/ece3.5664)

30. Roonwal ML. 1969 Measurements of termites for taxonomic purposes. J. Zool. Soc. India 21, 9–66.

31. Constantino R. 2016 Termite Database. Brasília, Univ. Brasília., updated Jul 2019.

32. Porter EE, Hawkins BA. 2001 Latitudinal gradients in colony size for social insects: Termites and ants show different patterns. Am. Nat. 157, 97–106. (doi:10.1086/317006)

33. Ronquist F, Klopfstein S, Vilhelmsen L, Schulmeister S, Murray DL, Rasnitsyn AP. 2012 A total-evidence approach to dating with fossils, applied to the early radiation of the hymenoptera. Syst. Biol. 61, 973–999. (doi:10.1093/sysbio/sys058)

34. Bourguignon T, Lo N, Šobotník J, Sillam-Dussès D, Roisin Y, Evans TA. 2016 Oceanic dispersal, vicariance and human introduction shaped the modern distribution of the termites Reticulitermes, Heterotermes and Coptotermes. Proc. R. Soc. B Biol. Sci. 283, 20160179. (doi:10.1098/rspb.2016.0179)

35. Bourguignon T et al. 2017 Mitochondrial phylogenomics resolves the global spread of higher termites, ecosystem engineers of the tropics. Mol. Biol. Evol. 34, 589–597. (doi:10.1093/molbev/msw253)

36. Bourguignon T et al. 2018 Transoceanic dispersal and plate tectonics shaped global cockroach distributions: Evidence from mitochondrial phylogenomics. Mol. Biol. Evol. 35, 970–983. (doi:10.1093/molbev/msy013)

37. Bucek A et al. 2021 Transoceanic voyages of ‘drywood’ termites (Isoptera: Kalotermitidae) inferred from extant and extinct species. bioRxiv (doi:10.1101/2021.09.24.461667)

38. Cameron SL, Whiting MF. 2007 Mitochondrial genomic comparisons of the subterranean termites from the genus Reticulitermes (Insecta: Isoptera: Rhinotermitidae). Genome 50, 188–202. (doi:10.1139/G06-148)

39. Scheffrahn RH, Bourguignon T, Akama PD, Sillam-Dussès D, Šobotník J. 2018 Roisinitermes ebogoensis gen. & sp. n., an outstanding drywood termite with snapping soldiers from Cameroon (isoptera, kalotermitidae). Zookeys 2018, 91–105. (doi:10.3897/zookeys.787.28195)

40. Wang M, Buček A, Šobotník J, Sillam-Dussès D, Evans TA, Roisin Y, Lo N, Bourguignon T. 2019 Historical biogeography of the termite clade Rhinotermitinae (Blattodea: Isoptera). Mol. Phylogenet. Evol. 132, 100–104. (doi:10.1016/j.ympev.2018.11.005)

41. Wu LW, Bourguignon T, Šobotník J, Wen P, Liang WR, Li HF. 2018 Phylogenetic position of the enigmatic termite family Stylotermitidae (Insecta: Blattodea). Invertebr. Syst. 32, 1111–1117. (doi:10.1071/IS17093)

42. Yamauchi MM, Miya MU, Nishida M. 2004 Use of a PCR-based approach for sequencing whole mitochondrial genomes of insects: Two examples (cockroach and dragonfly) based on the method developed for decapod crustaceans. Insect Mol. Biol. 13, 435–442. (doi:10.1111/j.0962-1075.2004.00505.x)

43. Ye F, Lan XE, Zhu WB, You P. 2016 Mitochondrial genomes of praying mantises (Dictyoptera, Mantodea): Rearrangement, duplication, and reassignment of tRNA genes. Sci. Rep. 6, 1–9. (doi:10.1038/srep25634)

44. Bernt M, Donath A, Jühling F, Externbrink F, Florentz C, Fritzsch G, Pütz J, Middendorf M, Stadler PF. 2013 MITOS: Improved de novo metazoan mitochondrial genome annotation. Mol. Phylogenet. Evol. 69, 313–319. (doi:10.1016/j.ympev.2012.08.023)

45. Katoh K, Standley DM. 2013 MAFFT multiple sequence alignment software version 7: Improvements in performance and usability. Mol. Biol. Evol. 30, 772–780. (doi:10.1093/molbev/mst010)

46. Suyama M, Torrents D, Bork P. 2006 PAL2NAL: Robust conversion of protein sequence alignments into the corresponding codon alignments. Nucleic Acids Res. 34, 609–612. (doi:10.1093/nar/gkl315)

47. Engel MS, Barden P, Riccio ML, Grimaldi DA. 2016 Morphologically specialized termite castes and advanced sociality in the early cretaceous. Curr. Biol. 26, 522–530. (doi:10.1016/j.cub.2015.12.061)

48. Lewis PO. 2001 A likelihood approach to estimating phylogeny from discrete morphological character data. Syst. Biol. 50, 913–925. (doi:10.1080/106351501753462876)

49. Harmon LJ, Weir JT, Brock CD, Glor RE, Challenger W. 2008 GEIGER: Investigating evolutionary radiations. Bioinformatics 24, 129–131. (doi:10.1093/bioinformatics/btm538)

50. Sallan LC, Friedman M. 2012 Heads or tails: Staged diversification in vertebrate evolutionary radiations. Proc. R. Soc. B Biol. Sci. 279, 2025–2032. (doi:10.1098/rspb.2011.2454)

51. Slater GJ, Harmon LJ, Alfaro ME. 2012 Integrating fossils with molecular phylogenies improves inference of trait evolution. Evolution (N. Y). 66, 3931–3944. (doi:10.1111/j.1558-5646.2012.01723.x)

52. R Core Team. 2020 R: A language and environment for statistical computing.

53. Orme D. 2018 The caper package: comparative analyses in phylogenetics and evolution in R.

54. Powell S, Price SL, Kronauer DJC. 2020 Trait evolution is reversible, repeatable, and decoupled in the soldier caste of turtle ants. Proc. Natl. Acad. Sci., 201913750. (doi:10.1073/pnas.1913750117)

55. Pequeno PACL, Baccaro FB, Souza JLP, Franklin E. 2017 Ecology shapes metabolic and life history scalings in termites. Ecol. Entomol. 42, 115–124. (doi:10.1111/een.12362)

56. Pequeno PACL, Graça MB, Oliveira JR, Šobotník J, Acioli ANS. 2020 Can shifts in metabolic scaling predict coevolution between diet quality and body size? Evolution (N. Y)., 1–8. (doi:10.1111/evo.14128)

57. Chown SL, Gaston KJ. 2010 Body size variation in insects: A macroecological perspective. Biol. Rev. 85, 139–169. (doi:10.1111/j.1469-185X.2009.00097.x)

58. Rainford JL, Hofreiter M, Mayhew PJ. 2016 Phylogenetic analyses suggest that diversification and body size evolution are independent in insects. BMC Evol. Biol. 16, 1–17. (doi:10.1186/s12862-015-0570-3)

59. Pagel M. 1999 Inferring historical patterns of biological evolution. Nature 401, 877–884.

60. Rabosky DL, Lovette IJ. 2008 Explosive evolutionary radiations: Decreasing speciation or increasing extinction through time? Evolution (N. Y). 62, 1866–1875. (doi:10.1111/j.1558-5646.2008.00409.x)

61. Nalepa CA, Maekawa K, Shimada K, Saito Y, Arellano C, Matsumoto T. 2008 Altricial development in subsocial wood-feeding cockroaches. Zoolog. Sci. 25, 1190–1198. (doi:10.2108/zsj.25.1190)

62. Nalepa CA. 2011 Altricial development in wood-feeding cockroaches: the key antecedent of termite eusociality. In Biology of Termites: A Modern Synthesis (eds DE Bignell, Y Roisin, N Lo), pp. 69–96. Springer. (doi:10.1007/978-90-481-3977-4)

63. Revell LJ. 2012 phytools: An R package for phylogenetic comparative biology (and other things). Methods Ecol. Evol. 3, 217–223. (doi:10.1111/j.2041-210X.2011.00169.x)

64. Mizumoto N, Gile GH, Pratt SC. 2020 Behavioral rules for soil excavation by colony founders and workers in termites. Ann. Entomol. Soc. Am., 1–8. (doi:10.1093/aesa/saaa017)

65. Korb J, Buschmann M, Schafberg S, Liebig J, Bagneres A-G. 2012 Brood care and social evolution in termites. Proc. R. Soc. London B 279, 2662–2671. (doi:10.1098/rspb.2011.2639)

66. Fjerdingstad EJ, Crozier RH. 2006 The evolution of worker caste diversity in social insects. Am. Nat. 167, 390–400. (doi:10.1086/499545)

67. Noirot C, Darlington JPEC. 2000 Termite nests: architecture, regulation and defence,. In Termites: Evolution, Sociality, Symbioses, Ecology (eds T Abe, DE Bignell, M Higashi), pp. 121–139. Netherlands: Kluwer Academic Publishers.

68. Prestwich GD. 1984 Defense mechanisms of termites. Annu. Rev. Entomol. 29, 201–232. (doi:10.1146/annurev.en.29.010184.001221)

69. Kalinkat G, Jochum M, Brose U, Dell AI. 2015 Body size and the behavioral ecology of insects: Linking individuals to ecological communities. Curr. Opin. Insect Sci. 9, 24–30. (doi:10.1016/j.cois.2015.04.017)

70. Bonner JT. 2011 Why size matters: From bacteria to blue whales. Princeton: Princeton University Press.

71. Ocko SA, Heyde A, Mahadevan L. 2019 Morphogenesis of termite mounds. Proc. Natl. Acad. Sci. U. S. A. 116, 3379–3384. (doi:10.1073/pnas.1818759116)

72. Mizumoto N, Bardunias PM, Pratt SC. 2020 Complex relationship between tunneling patterns and individual behaviors in termites. Am. Nat. 196, 555–565. (doi:10.1086/711020)

73. Theraulaz G, Bonabeau E. 1995 Coordination in distributed building. Science (80-.). 269, 686–688. (doi:10.1126/science.269.5224.686)

74. Camazine S, Deneubourg J-L, Franks NR, Sneyd J, Theraulaz G, Bonabeau E. 2001 Self-organization in Biological Systems. Princeton: NJ: Princeton University Press.

75. Khuong A, Theraulaz G, Jost C, Perna A, Gautrais J. 2011 A computational model of ant nest morphogenesis. Adv. Artif. Life, ECAL 2011, Proc. Elev. Eur. Conf. Synth. Simul. Living Syst., 404–411.

